# Earlier springs enable High-Arctic wolf spiders to produce a second clutch

**DOI:** 10.1101/2020.04.30.070029

**Authors:** Toke T. Høye, Jean-Claude Kresse, Amanda M. Koltz, Joseph J. Bowden

## Abstract

Spiders at southern latitudes commonly produce multiple clutches, but this has not been observed at high latitudes where activity seasons are much shorter. Yet the timing of snowmelt is advancing in the Arctic, and may allow some species to produce an additional clutch. To determine if this is already happening, we used specimens of the wolf spider *Pardosa glacialis* caught by pitfall traps from the long-term (1996-2014) monitoring program at Zackenberg, Northeast Greenland. We dissected individual egg sacs and counted the number of eggs and partially developed juveniles, and measured carapace width of the mothers. Upon discovery of a bimodal frequency distribution of clutch sizes, as is typical for wolf spiders at lower latitudes producing a second clutch, we assigned egg sacs to being a first or second clutch depending on clutch size. We tested whether the median capture date differed among first and second clutches, whether clutch size was correlated to female size, and whether the proportion of second clutches produced within a season was related to climate. We found that assigned second clutches appeared significantly later in the season than first clutches. In years with earlier snowmelt, first clutches occurred earlier and the proportion of second clutches produced was larger. This result, likely a result of female spiders producing first clutches earlier in those years, which allowed time for another clutch. Clutch size for first clutches was correlated to female size, while this was not the case for second clutches. Our results provide the first evidence for Arctic invertebrates producing additional clutches in response to warming. This could be a common but overlooked phenomenon due to the challenges associated with long-term collection of life history data in the Arctic. Moreover, given that wolf spiders are a widely distributed, important tundra predator, we may expect to see population and food web consequences of their increased reproductive rates.

## INTRODUCTION

Shifts in phenology, i.e. the timing of biological events, are the most widely reported biological response to climate change [1–3]. The demographic consequences of climate-induced phenological variation are typically studied in the context of phenological mismatch, where mistiming of resource availability and resource demand lead to reduced growth, survival, or reproduction [4–6]. Some studies have examined how the number of generations per year may increase in response to a warmer climate [7–9], but little is known about how extended growing seasons may affect total reproductive output [10, 11]. Yet warmer conditions can extend the time available for reproduction, which may result in phenological shifts that could also affect demography and population dynamics [12]. Due to the low temperatures and time limitations of short growing seasons in northern environments, invertebrate organisms that typically produce multiple clutches in temperate ecosystems can typically only produce one clutch at higher latitudes [13, 14]. Arctic temperatures are currently rising at twice the global average and climate projections indicate that the Arctic will continue to warm at a higher rate than the rest of the globe [15]. These warmer conditions and associated extended growing seasons may result in higher reproductive rates among Arctic species that are opportunistically capable of producing additional clutches.

Arctic arthropods are expected to be particularly affected by climate, and changes in their phenology have been widely documented [16–18]. Wolf spiders (family: Lycosidae) are abundant on the Arctic tundra [19–23] and they play an important role as top predators in the ecosystem [24–26]. They are also very responsive to environmental change [27, 28]. In the North, their life cycle typically takes two or more years [29, 30], while in the temperate zone, wolf spiders have annual life cycles [31]. Adult spiders die after completing reproduction, but female wolf spiders at lower latitudes typically produce more than one clutch over their lifetimes [32, 33]. Subsequent egg clutches are produced approximately one month after the first clutch, and they differ from first clutches in that there are typically fewer eggs produced per clutch [32]. In the Arctic, due to the time constraints of extremely short growing seasons, female wolf spiders have always been assumed to produce only one clutch [34]. However, the opportunity to produce a second clutch could confer a big fitness advantage, which may be possible with the warmer temperatures and extended growing seasons that are currently being brought on by climate change [35]. The characteristics associated with the production of second clutches in temperate regions suggest that if longer growing seasons enable female wolf spiders in the Arctic to produce second clutches, these clutches would likely occur later in the summer, contain fewer eggs per clutch, and occur more frequently in years with early snowmelt or higher temperatures.

Longer growing seasons may also increase reproductive rates indirectly through changes in female body size. Fecundity is typically positively associated with body size in invertebrates [36], including among spiders, whereby larger females produce larger clutches [37, 38]. Body size – and hence fecundity – also vary according to growing season length in these organisms. For example, previous studies from multiple Arctic locations have found that wolf spider body size and fecundity decrease with rising elevation, a proxy for shortening growing season length [39] [40]. Whether larger females are also be more likely to produce second clutches is unknown. We would expect body size of females producing a second clutch to be larger if only the biggest individuals are able to produce two clutches, or alternatively, females should be of similar size if the ability to produce two clutches mainly depends on the length of the season.

Here, we use long-term (1996–2014) data on clutch size variation in the only wolf spider species *(Pardosa glacialis,* Thorell 1872) known from the study area at Zackenberg, NE Greenland to examine if climate change is enabling the spiders to produce an additional clutch through a lengthening of the growing season. Previous work has found that body sizes of wolf spiders at this High-Arctic site are larger in years with earlier snowmelt [41]. Climate change at Zackenberg in NE Greenland is resulting in warmer summer temperatures and earlier spring snowmelt/growing season [42]. We argue that the combination of three patterns would indicate that second clutches are produced in a population: 1) A bimodal frequency distribution of clutch sizes which would allow for an assignment of clutches to either first or second clutches. 2) Females with egg sacs assigned to first clutches should be caught earlier than those females with egg sacs assigned to second clutches. 3) Females producing clutches assigned as first clutches should not be bigger than females carrying egg sacs that are assigned as second clutches. We use data from a long-term (1996-2014) standardized monitoring program at Zackenberg in High-Arctic Greenland to examine the likelihood of whether second clutches are being produced by female wolf spiders. We make the following predictions in light of climate change: 1) A proportion of all egg sacs are from second clutches, 2) that this proportion varies across years at the site, and 3) the proportion of second clutches is related to timing of snowmelt date.

## MATERIALS AND METHODS

### Field and lab work

Female wolf spiders carry their egg sac attached to their spinnerets, and therefore pitfall trap samples collected across the breeding season can be used to assess the prevalence of potential second clutches. We used samples of wolf spiders that were collected from pitfall traps in five different plots at Zackenberg, Northeast Greenland (74°28N, 20°34’W, 35-50 m.a.s.l.) as part of the Greenland Ecosystem Monitoring program. The plots are placed in three different habitats with one plot in a fen and two plots each in mesic heath and arid heath habitats. Pitfall traps were emptied weekly during June, July and August, although trapping continued into September in a few years. In order to standardize trapping effort, we omitted samples (totalling 17 egg sacs) collected before 1 June and after 27 August. Only one species of wolf spider *(P. glacialis)* has been collected from the study area over this 18 year period [27], so all egg sacs are assumed to belong to this species. The species has a generation time of two years at the site [41]. All egg sacs from years 1996–2014 (no available data for 2010) were opened. We counted the number of eggs or partially developed juveniles. If the egg sac was still attached to the mother, we measured the width of her carapace to the nearest 0.01 mm using a reticle fitted to a stereo microscope. Otherwise, the mother could only be identified if there was only one female in the sample. A total of 1069 egg sacs were collected across the 18 years of sampling. The mother could only be confidently identified in 280 of these cases due to egg sacs detaching from the mother during trapping. We calculated the date of snowmelt across plots as the average date at which 50% of the snow had disappeared from each plot during spring snowmelt, measured as the day of the year after 1 January.

### Data analysis

We found a clear bimodal frequency distribution of the sizes of clutches and could therefore assign each clutch to a first or second clutch depending on clutch size. We tested if there was a significant difference in the mean capture dates between egg sacs that were assigned to first or second clutches using a one-way ANOVA. We then tested whether the proportion of second clutches produced across years varied among habitat types. Likewise, we tested whether this proportion exhibited a trend across the study period or was related to the date of snowmelt using general linear models with a Gaussian error distribution. We also tested if the mean date of trapping first clutches was related to timing of snowmelt. Finally, for the subset of data for which the mother could be identified, we tested if clutch size was related to body size of the mother separately for first and second clutches. For this subset, we also tested if mothers that were assigned to first or second clutches differed in body size using a general linear model. All analyses were conducted in R (R Core Development Team 2016).

## RESULTS

Clutch sizes exhibited a distinct bimodal distribution with a split between peaks of clutch size at 47 eggs (Figure 1). We assigned clutches with more than 47 eggs to first clutches and considered clutches with 47 eggs or less to be second clutches. Clutches with 47 eggs or less were caught significantly later in the season than larger clutches (difference = 20.7±1.3 days, F_1,1067_ = 253.6, p<0.001; Figure 2). The same was true within individual years based on comparisons of the average dates of at least ten egg sacs in first and second clutches (data not shown). Of the 17 omitted egg sacs collected after 27 August, 15 (88%) had 47 eggs or less and, therefore, would have been assigned to second clutches. Across all years, the proportion of second clutches varied among habitats and was highest in the wet fen (arid heath = 0.113, mesic heath = 0.128, and wet fen = 0.543).

**Figure 1.**
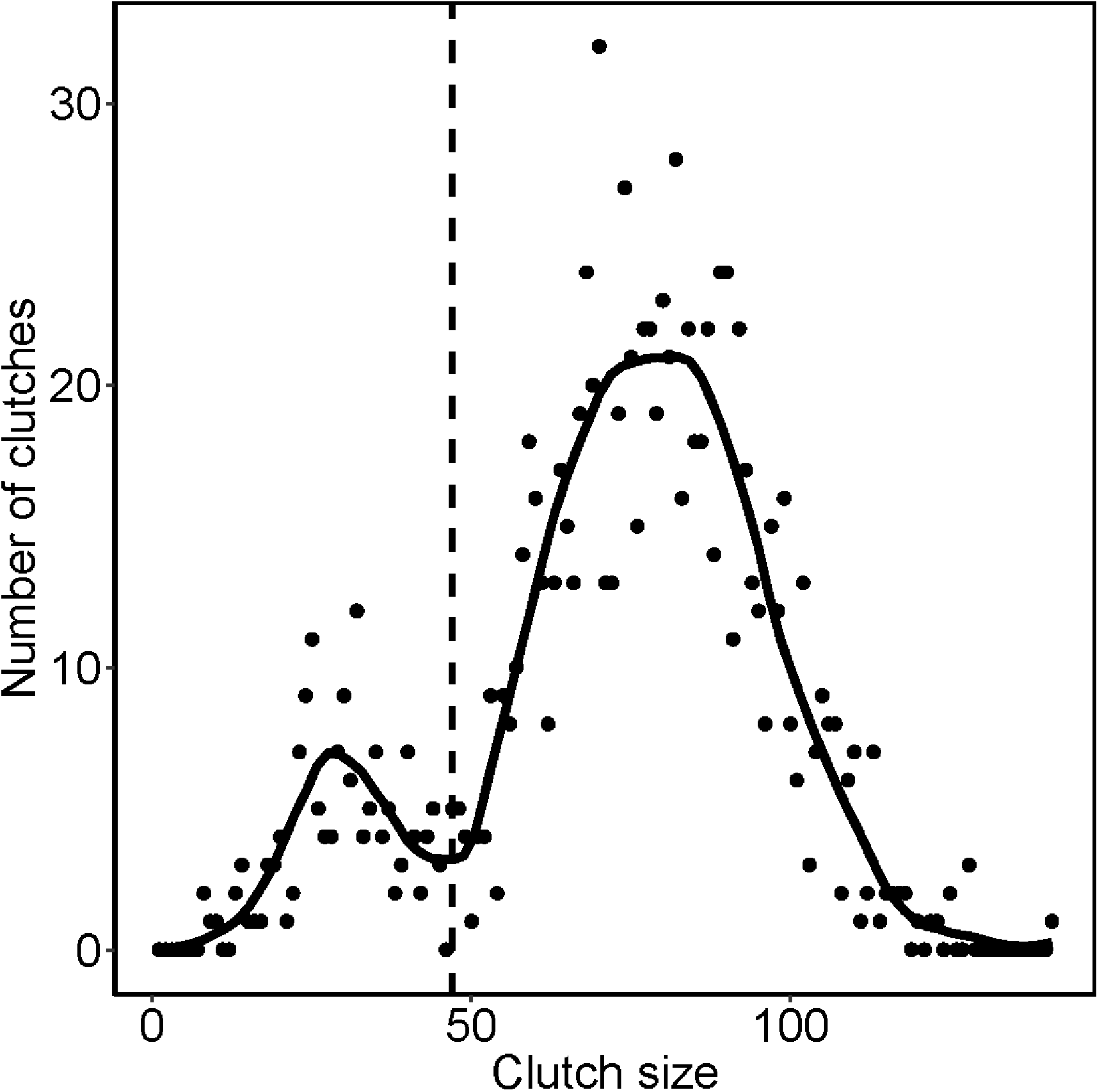
Frequency distribution of clutch sizes in the wolf spider *Pardosa glacialis* across 1069 egg sacs collected in pitfall traps during 1996-2014 at Zackenberg, North-East Greenland. The solid line represents a locally weighted smoothing with a span parameter = 0.2 and identifies a local minimum at a clutch size of 47 eggs used to separate first and second clutches, as indicated by the vertical hatched line.

**Figure 2.**
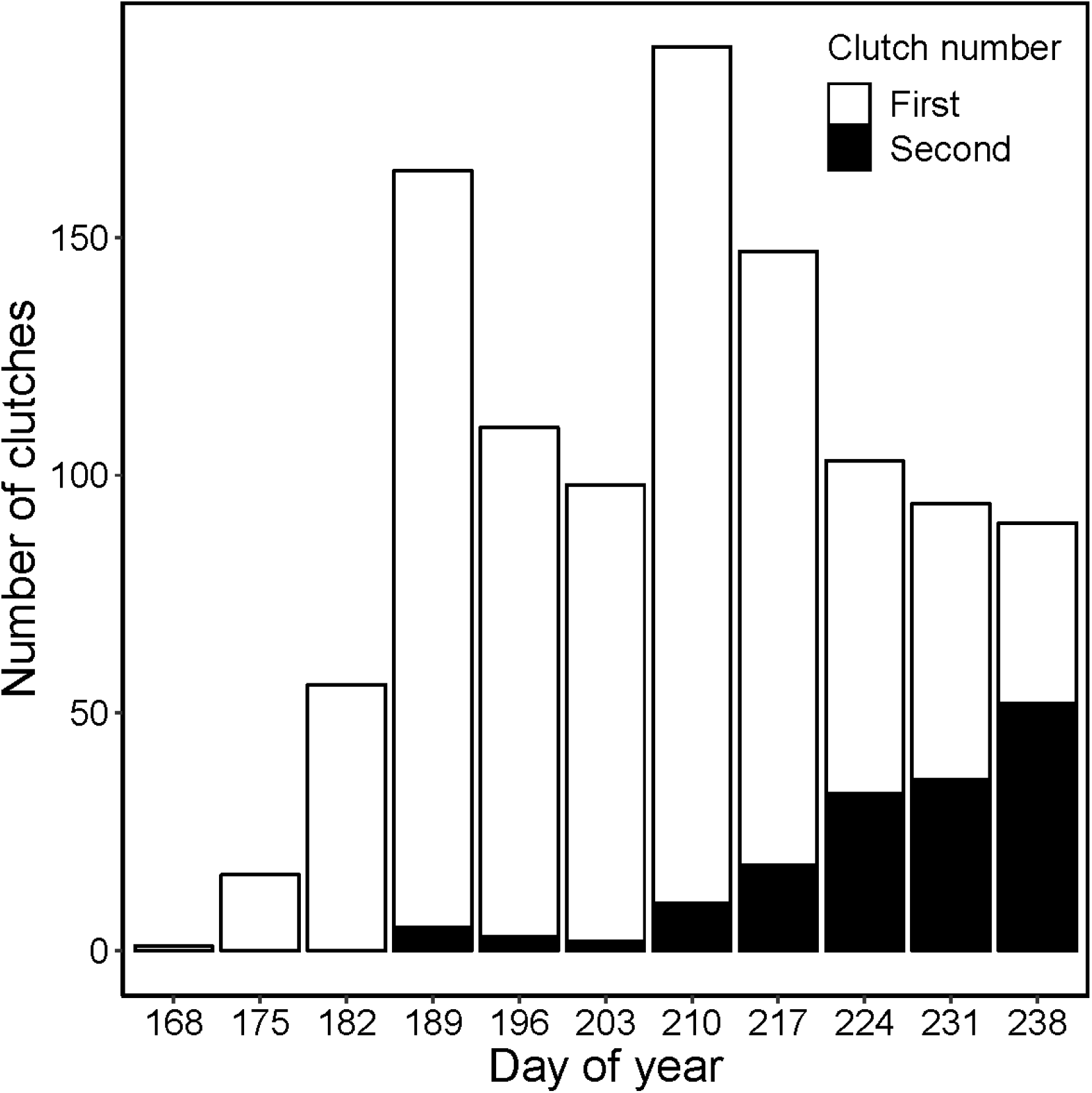
Seasonal variation in total egg sac collection across the study period (grouped by seven day periods). Dates are presented as day of year since 1 January. The white section of each bar indicates clutches with more than 47 eggs (first clutches) and the black sections are for those clutches with 47 or less eggs (second clutches).

Snowmelt advanced significantly during the study period (slope = −0.79±0.36, R^2^_adj_ = 0.18, n = 18, p = 0.043). We found a negative relationship between the proportion of second clutches and date of snowmelt (slope = −0.015±0.0038, R^2^_adj_ = 0.46, n = 18, p = 0.0012; Figure 3a) and sampling year (slope = 0.0182±0.0075, R^2^_adj_ = 0.22, n = 18, p = 0.028; Figure 3b). We also found earlier mean capture date of first clutches in years with earlier snowmelt (slope = 0.90±0.20, R^2^_adj_ = 0.54, n = 18, p = 0.0032). Clutch size of first clutches was largest in wet habitats, intermediate in mesic habitats and smallest in arid habitats (arid vs. wet = 28.40±3.29, t = 8.65, p < 0.001; arid vs. mesic = 5.66±1.56, t = 3.63, p = 0.00035) and was positively related to carapace width of the mother (estimate = 68.48±5.15, t = 13.29, p < 0.0001; Figure 4a). The size of second clutches was independent of habitats (arid vs. mesic = 0.60±3.93, t = 0.15, p = 0.88; arid vs. wet = 2.45±4.10, t = 0.60, p = 0.56) and unrelated to carapace width of mothers (estimate = 3.10±8.2, t = 0. 38, p = 0.71; Figure 4b). There was no significant difference in body size between mothers of first and second clutches (difference = −0.0039±0.025, F1,278 = 0.024, p = 0.88).

**Figure 3.**
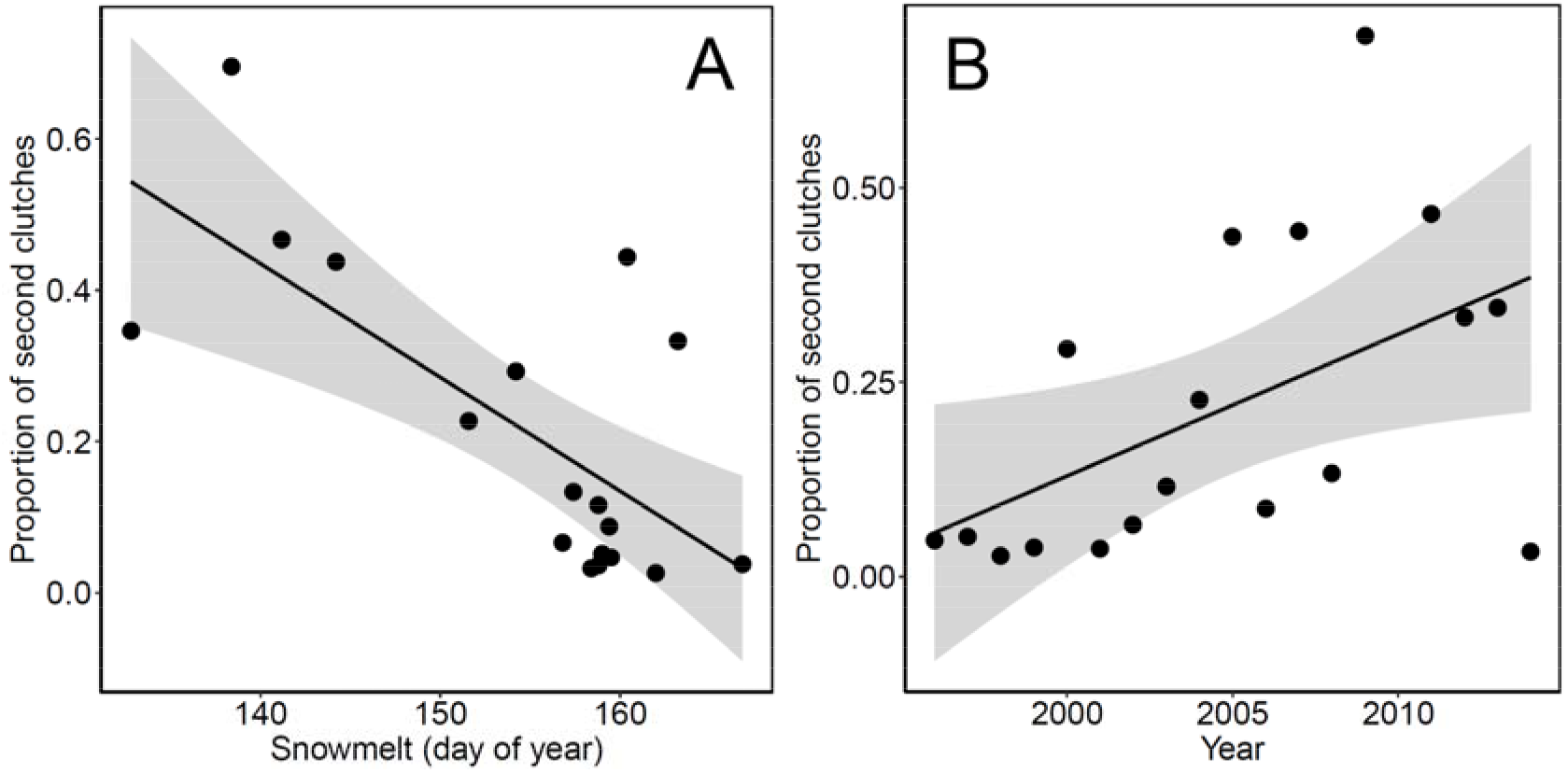
The proportion of second clutches out of total number of clutches in a given year regressed against a) average date of snowmelt (day of year since January 1) across the study plots (slope = – 0.0150±0.0038, t_16_ = −3.949, p = 0.0012) and b) sampling year (slope = 0.0182±0.0075, t_16_ = 2.418, p = 0.028).

**Figure 4.**
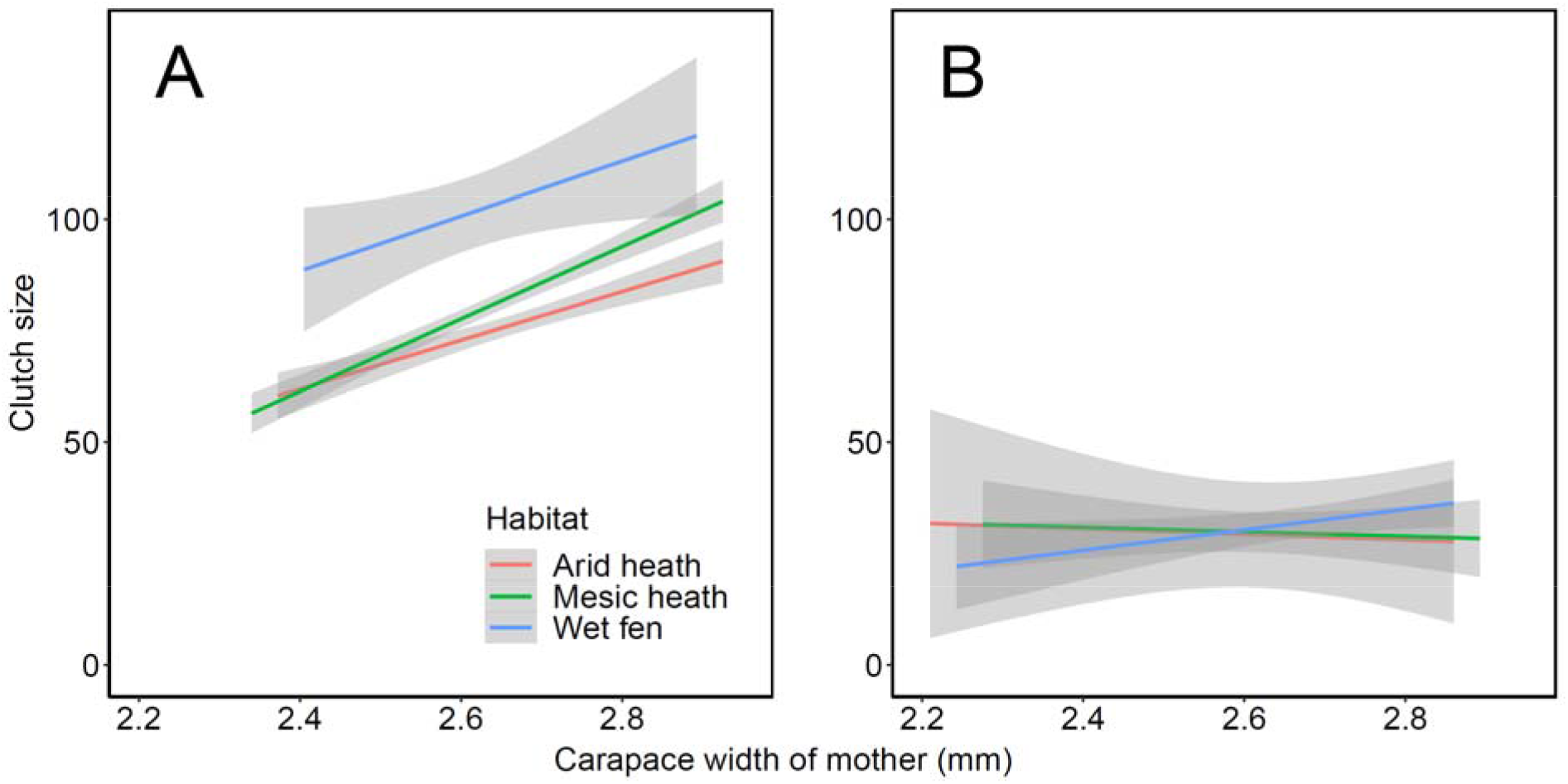
Relationship between clutch size and body size of the mother for each habitat type (wet, mesic, and arid) for (A) first and (B) second clutches. The relationships are significant for first clutches, but not for second clutches (see text for details). Body size of the mother was estimated as width of the carapace (in millimeters).

## DISCUSSION

Over the last two decades, the climate at Zackenberg has changed significantly, including snowmelt occurring progressively earlier and temperatures becoming warmer. These changes have had rapid measurable impacts on the biology of species in the area [43, 44]. In addition to previously documented changes, we have detected strong evidence of climate change induced shifts to multivoltinism in a dominant spider species. Our results provide the first evidence of an arctic spider species being able to produce two clutches. This is demonstrated by a striking bimodal frequency distribution of clutch sizes in the wolf spider *P. glacialis*, which is indicative of second clutches in temperate regions [32, 45]. Moreover, clutches with fewer eggs (47 eggs or less) were produced significantly later in the season than larger clutches. We are able to rule out the possibility that this was driven by inter-annual variation in egg sac phenology, because second clutches were laid later in the season in all years. Together, these findings provide support for the hypothesis that *P. glacialis* is able to produce a second clutch at our high-arctic study site in North-East Greenland. We were further able to demonstrate that as spring snowmelt becomes earlier, a greater proportion of *P. glacialis* females are able to produce a second clutch and that the production of second clutches is increasing over time. In fact, the proportion of second clutches increased from zero to >50% as snowmelt became progressively earlier over the study period, suggesting that the ability to produce second clutches is a common yet overlooked phenomenon in northern ecosystems.

It is known that larger female wolf spiders produce more offspring [46] and our results confirm this for first clutches. However, we found no indication that larger females are more likely to produce a second clutch than smaller females or that larger females produce larger second clutches. These findings suggest that the production of second clutches in association with earlier snowmelt is not due to females obtaining more resources and attaining larger body sizes. Rather, earlier snowmelt enables females to produce their first clutch earlier and gives them enough time to produce a second clutch before the season ends. It is possible that climate change is altering the reproductive strategy of *P. glacialis*, allowing females to produce second clutches for the first time. However, given that Arctic wolf spiders have been shown to respond rapidly to environmental conditions and given the broad distribution of individual species [34], it seems plausible that this species is already adapted to opportunistically reproduce a second time when conditions allow. Although our study was limited to a single site, Zackenberg is at the northern edge of the range of *P. glacialis* and one of the most climatically extreme locations inhabited by this species. Thus, it is likely that *P. glacialis* and possibly other wolf spider species are also already producing second clutches at lower arctic and boreal latitudes. Similar studies at other sites and for other invertebrate species are needed to unravel the ubiquity of climate-induced shifts in voltinism and the potential ecological consequences.

We found a difference in the mean date of capture of first and second clutches of about 20 days. This period is less than previous reports of a 30-day interval between the production of first and second clutches in temperate wolf spiders [32], but it may be related to the long daily hours of incident solar radiation at high latitudes. The growing season at Zackenberg is extremely short and *P. glacialis* could be adapted to rapidly respond to favorable conditions by producing second clutches whenever opportunity allows. It is possible that some of the clutches that were assigned as second clutches could have been from females who lost their first egg sacs (e.g. due to predation from birds). In such cases, the production of a second, smaller clutch would happen earlier in the season and would reduce our estimate of a time difference. Still, the combination of evidence presented here strongly suggest that *P. glacialis* are indeed producing second clutches in years with earlier snowmelt at our High-Arctic site.

Increased reproductive output can increase population size if other factors such as competition or predation are not limiting population growth. Yet, we have not observed significant changes in population size of *P. glacialis* at this site during the study period [27]. Thus, it is unclear whether increased reproductive rates associated with the production of second clutches will have population-level consequences for this species in the future. Recent evidence from our field site suggests that *P. glacialis* is not food limited [47], nor are its populations affected by parasitism as in other arctic wolf spiders [48, 49]. However, wolf spiders are density-dependent cannibals [e.g., 50, 51, 52], and a lack of observed population growth could be a result of increased intraspecific competition with rising reproductive rates [53]. As *P. glacialis* is the largest invertebrate predator at our site, further studies should address whether there are implications of higher reproductive rates among the spiders for prey populations. If prey populations do not respond in a similar way to earlier snowmelt, they might be negatively affected by an increasing predation pressure as top-down forces appears to be the main control mechanism of tundra food webs [26, 54, 55]. Smaller arthropod predators, such as other spiders, might also suffer from the increase in abundance of *P. glacialis* due to the limited differentiation in diet between predators in the High Arctic [27, 53, 56]. Studies of the effects on population dynamics, cascading effects in food webs, and repercussions on ecosystem function, continue to establish wolf spiders as key model organisms for the study of Arctic climate change effects.

## ACKNOWLEDGEMENTS

Access to spider specimens and climate data from the Greenland Ecosystem Monitoring program is greatly appreciated. Spider specimens are curated by the Natural History Museum, Aarhus, Denmark.

